# Sentinel2GlobalLULC: A deep-learning-ready Sentinel-2 RGB image dataset for global land use/cover mapping

**DOI:** 10.1101/2021.12.01.470768

**Authors:** Yassir Benhammou, Domingo Alcaraz-Segura, Emilio Guirado, Rohaifa Khaldi, Boujemâa Achchab, Francisco Herrera, Siham Tabik

## Abstract

Land-Use and Land-Cover (LULC) mapping is relevant for many applications, from Earth system and climate modelling to territorial and urban planning. Global LULC products are continuously developing as remote sensing data and methods grow. However, there is still low consistency among LULC products due to low accuracy for some regions and LULC types. Here, we introduce Sentinel2GlobalLULC, a Sentinel-2 RGB image dataset, built from the consensus of 15 global LULC maps available in Google Earth Engine. Sentinel2GlobalLULC v1.1 contains 195572 RGB images organized into 29 global LULC mapping classes. Each image is a tile that has 224 × 224 pixels at 10 × 10 m spatial resolution and was built as a cloud-free composite from all Sentinel-2 images acquired between June 2015 and October 2020. Metadata includes a unique LULC type annotation per image, together with level of consensus, reverse geo-referencing, and global human modification index. Sentinel2GlobalLULC is optimized for the state-of-the-art Deep Learning models to provide a new gate towards building precise and robust global or regional LULC maps.

## 1 Background & Summary

Land-Use and Land-Cover mapping aims to comprise the continuous biophysical properties of the Earth surface into synthetic categorical classes of natural or human origin, such as forests, shrublands, grasslands, marshlands, croplands, urban areas or water bodies^1^. High resolution LULC mapping plays a key role in many fields, from natural resources monitoring, to biodiversity conservation, urban planning, agricultural management or climate and earth system modelling^2–4^. Multiple LULC products have been derived using satellite information at the global scale (Table 2), contributing to a better monitoring and understanding of our planet^5,6^. However, despite the acceptable accuracy of each individual product, a considerable disagreement between products has been reported^4,7–22^. There are several methodological reasons behind this problem:

- Different satellite sensors with different spatial resolutions were used in each product, so the difference in precision from coarse to fine resolution partially determines the final quality of each product.
- Different pre-processing techniques, like atmospheric corrections, cloud removal and image composition were used in each LULC product.
- Each LULC product has a different temporal updating rate, some are regularly updated, whereas others have never been updated.
- Different classification techniques, field-data collection approaches and subjective interpretations were used to create each product.
- Different classification systems (LULC legends) were adopted in each product, usually focused on distinct applications.
- Different validation techniques and different ground truth reference data were used in each product, which impedes a reliable accuracy comparison.

Over the last few years, several attempts have been made to overcome these inconsistencies with a harmonised approach capable of providing greater control in the validation and comparison over the growing number of existing LULC products^23,24^. Even though, users still have some issues regarding appropriate product selection due to the following factors:

- In most cases, users are unable to find a product that fits their desired LULC class or geographic region of interest^25,26^.
- These products are usually collected at a coarse resolution, which makes analysis at a finer scale difficult^12^.
- These products offers a limited number of LULC classes that usually change from one product to another^27^.

In parallel, Deep artificial neural networks, also known as Deep learning (DL), are increasingly used in LULC mapping with promising potential^28^. This interest is motivated by the good performance of DL models in computer vision and, particularly of Convolutional Neural Networks (CNNs) in remote sensing image classification and many applications^29–33^. However, to reach high performance, DL models need to be trained on large smart datasets^34^. The concept of smart data involves all pre-processing methods that improve value and veracity of the data in addition to the quality of the associated expert annotations^35^.

Currently, there exist several remote sensing datasets derived from satellite and aerial imagery ready for training DL models for LULC mapping (Table 1). However, they still suffer from some limitations, particularly to be used with DL models:

**Table 1.**
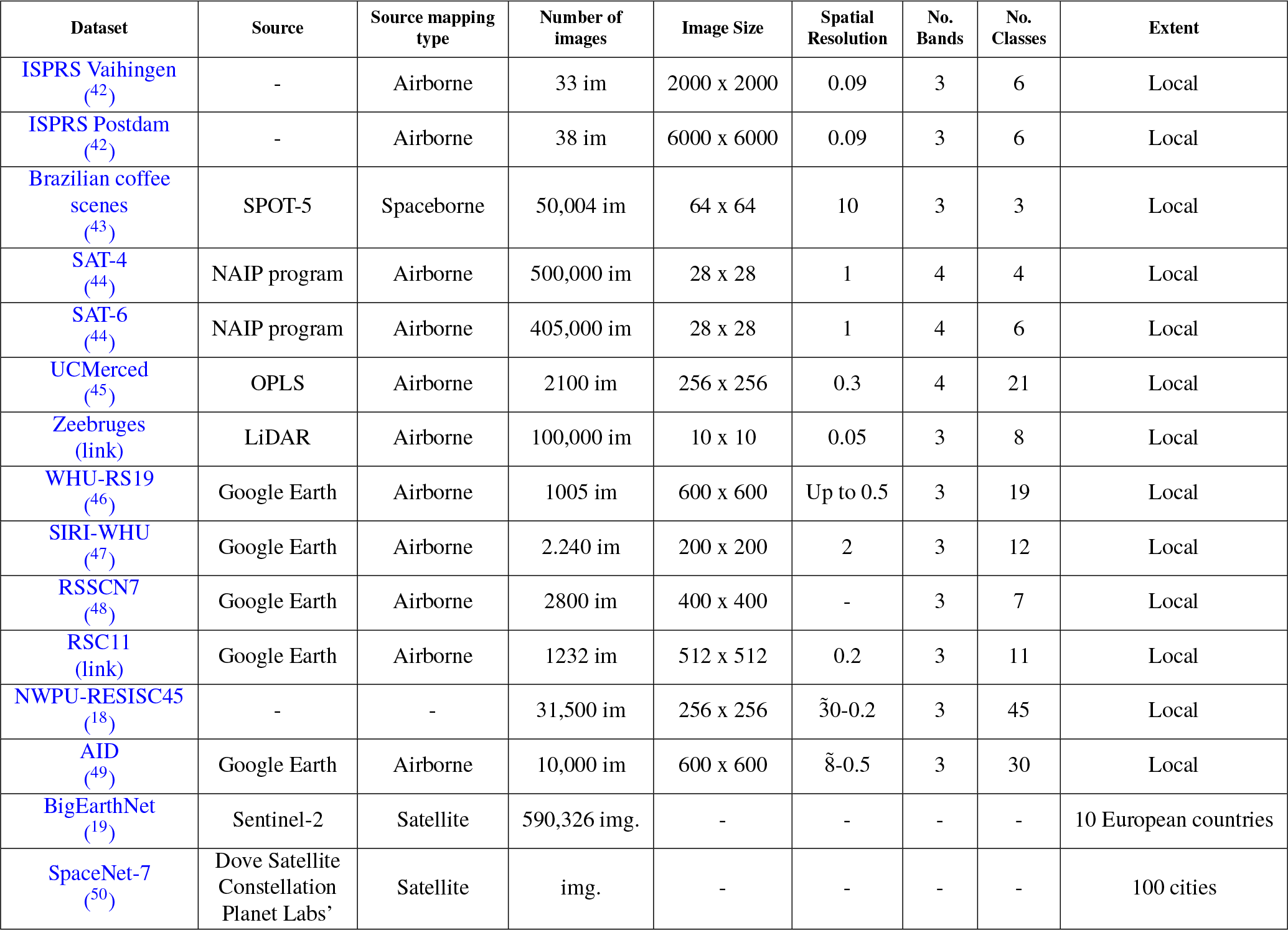
List of existing Land-Use and Land-Cover (LULC) datasets ready for training Deep Learning (DL) models.

| Dataset | Source | Source mapping type | Number of images | Image Size | Spatial Resolution | No. Bands | No. Classes | Extent |
| --- | --- | --- | --- | --- | --- | --- | --- | --- |
| ISPRS Vaihingen<sup>(42)</sup> | - | Airborne | 33 im | 2000 x 2000 | 0.09 | 3 | 6 | Local |
| ISPRS Postdam<sup>(42)</sup> | - | Airborne | 38 im | 6000 x 6000 | 0.09 | 3 | 6 | Local |
| Brazilian coffee scenes<sup>(43)</sup> | SPOT-5 | Spaceborne | 50,004 im | 64 x 64 | 10 | 3 | 3 | Local |
| SAT-4<sup>(44)</sup> | NAIP program | Airborne | 500,000 im | 28 x 28 | 1 | 4 | 4 | Local |
| SAT-6<sup>(44)</sup> | NAIP program | Airborne | 405,000 im | 28 x 28 | 1 | 4 | 6 | Local |
| UCMerced<sup>(45)</sup> | OPLS | Airborne | 2100 im | 256 x 256 | 0.3 | 4 | 21 | Local |
| Zeebruges (link) | LiDAR | Airborne | 100,000 im | 10 x 10 | 0.05 | 3 | 8 | Local |
| WHU-RS19<sup>(46)</sup> | Google Earth | Airborne | 1005 im | 600 x 600 | Up to 0.5 | 3 | 19 | Local |
| SIRI-WHU<sup>(47)</sup> | Google Earth | Airborne | 2,240 im | 200 x 200 | 2 | 3 | 12 | Local |
| RSSCN7<sup>(48)</sup> | Google Earth | Airborne | 2800 im | 400 x 400 | - | 3 | 7 | Local |
| RSC11 (link) | Google Earth | Airborne | 1232 im | 512 x 512 | 0.2 | 3 | 11 | Local |
| NWPU-RESISC45<sup>(18)</sup> | - | - | 31,500 im | 256 x 256 | 30-0.2 | 3 | 45 | Local |
| AID<sup>(49)</sup> | Google Earth | Airborne | 10,000 im | 600 x 600 | 8-0.5 | 3 | 30 | Local |
| BigEarthNet<sup>(19)</sup> | Sentinel-2 | Satellite | 590,326 img. | - | - | - | - | 10 European countries |
| SpaceNet-7<sup>(50)</sup> | Dove Satellite Constellation Planet Labs' | Satellite | img. | - | - | - | - | 100 cities |

- None of them represent the global heterogeneity of the broad categories of LULC classes throughout the Earth. Usually, they are biased towards specific regions of the world, limited to national or continental scales, which can propagate such bias to the DL models^36–38^. As illustration, the reader can see how visual features of urban areas may change from one country to another (Figure 1).
- They are relatively small and have only hundreds to few thousands of annotated data records^39^.
- They suffer from high variability in atmospheric conditions, and they have high inter-class similarity and intra-class variability, which makes class differentiation difficult^39,39^.

To overcome these limitations, we introduce in this paper Sentinel2GlobalLULC, a smart dataset with 29 fully annotated LULC classes at global scale built with Sentinel-2 RGB imagery. Every image in this dataset is geo-referenced and labeled with its corresponding LULC annotation. Each image label was carefully built from a consensus approach of up to 15 global LULC maps available on [Google Earth Engine(GEE)](https://earthengine.google.com/)^40^. We released a tif and jpeg version of each image. Moreover, we attached to these images, a CSV file for each LULC class containing the coordinates of each image center, and additional metadata. Sentinel2GlobalLULC could be used to train and/or evaluate DL based models for global LULC mapping. Sentinel2GlobalLULC aims to foster the creation of accurate global LULC products exploiting the advantages that currently offer deep learning models. We expect this dataset to improve our understanding and modelling of natural and human systems around the world.

**Figure 1.**
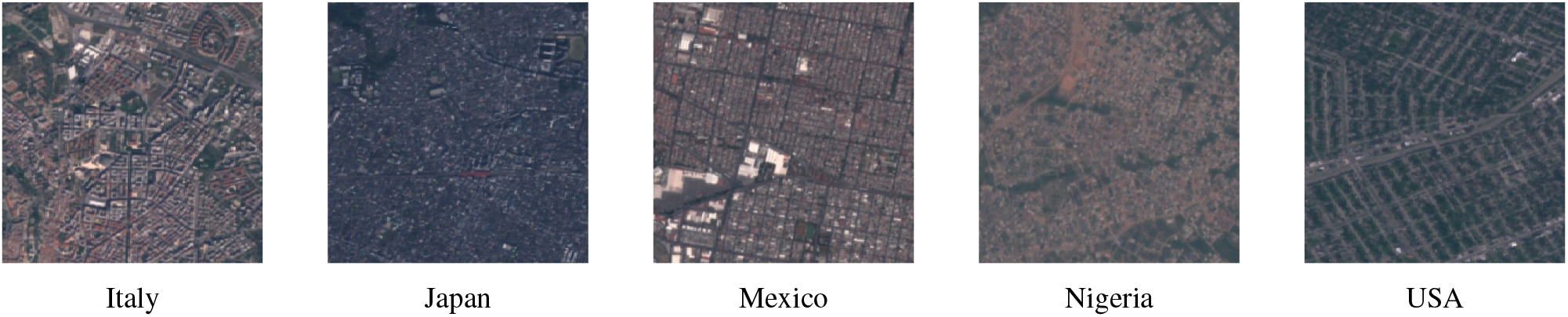
Illustration from different countries of the Sentinel-2 satellite images corresponding to one of the 29 Land-Use and Land-Cover (LULC) classes (e.g. Urban and built-up area) extracted from Sentinel2GlobalLULC dataset. Each image has 224 × 224 pixels of 10 × 10 m resolution. Pixel values were calculated as the 25th-percentile of all images captured between June 2015 and October 2020 that were not tagged as cloudy. Fifteen LULC products available in Google Earth Engine agreed in annotating each image to represent one LULC class

## 2 Methods

To build Sentinel2GlobalLULC, we followed two main steps. First, we established a spatial consensus between 15 global LULC products for 29 LULC classes. Then, for each class, we carefully extracted the maximum possible number of Sentinel-2 RGB images in 224 × 224 pixel tiles at 10 m/pixel spatial resolution. Both tasks were implemented using GEE, an efficient programming, processing and visualisation platform that allowed us to have free manipulation and access to all used LULC products and Sentinel-2 imagery, simultaneously.

### 2.1 Finding spatio-temporal agreement across 15 global LULC products

To establish the spatio-temporal consensus between different LULC products for each one of the 29 LULC classes, we followed four steps: 1) identification of the LULC products to use for the consensus, 2) standardization and harmonization of the LULC legend that was subsequently used as annotation, 3) spatio-temporal aggregation across selected LULC products, and 4) spatial reprojection and tile selection based on optimized spatial purity thresholds.

#### 2.1.1 Global LULC products selection

To find areas of high consensus in their LULC mapping, we selected the 15 global LULC products available in GEE (Table 2). Reaching consensus across such rich diversity of LULC products, in terms of spatial resolution, time coverage, satellite source, LULC classes and accuracy, made our LULC annotation robust. This way, each image was annotated with a LULC class only if all combined products agreed (i.e., 100% of agreement in space and time). For some LULC classes, we had to decrease the purity threshold to reach a large number of samples. The purity level is always provided as metadata for each image (details in the subsection Re-projection and Selection of purity threshold).

**Table 2.** Main characteristics of the 15 global Land-Use and Land-Cover (LULC) products available in Google Earth Engine (GEE) that were combined to find consensus in the global distribution of 29 main LULC classes

| LULC product | Satellite or Spaceborne | Resolution | Used years | Reference |
| --- | --- | --- | --- | --- |
| **P1:** MCD12Q1.006 MODIS LULC<br>Type Yearly Global 500m<br>LULC Type1: Annual International Geosphere-Biosphere Programme (IGBP) classification (version 6) | Aqua, Terra | 500 meters | 2017 to 2019 | <sup>51</sup> |
| **P2:** MCD12Q1.006 MODIS LULC<br>Type Yearly Global 500m<br>LULC Type 2: Annual University of Maryland (UMD) classification (version 6) | Aqua, Terra | 500 meters | 2017 to 2019 | <sup>51</sup> |
| **P3:** MCD12Q1.006 MODIS LULC<br>Type Yearly Global 500m<br>LULC Type 3: Annual Leaf Area Index (LAI) classification (version 6) | Aqua, Terra | 500 meters | 2017 to 2019 | <sup>51</sup> |
| **P4:** MCD12Q1.006 MODIS LULC<br>Type Yearly Global 500m<br>LULC Type 4: Annual BIOME-Biogeochemical Cycles (BGC) classification (version 6) | Aqua, Terra | 500 meters | 2017 to 2019 | <sup>51</sup> |
| **P5:** MCD12Q1.006 MODIS LULC<br>Type Yearly Global 500m<br>LULC Type 5: Annual Plant Functional Types classification (version 6) | Aqua, Terra | 500 meters | 2017 to 2019 | <sup>51</sup> |
| **P6:** Copernicus Global LULC Layers: CGLS-LC100 collection 3 (version 3.0.1) | PROBA-V | 100 meters | 2017 to 2019 | <sup>52</sup> |
| **P7:** Global Forest Cover Change (GFCC) Tree Cover Multi-Year Global 30m (version 3.0) | Multi-satellite | 30 meters | 2015 | <sup>53</sup> |
| **P8:** GlobCover: Global LULC Map (version 2.0) | ENVISAT | 300 meters | 2009 | ESA 2010 and UCLouvain |
| **P9:** GFSAD1000: Cropland Extent 1km Multi-Study Crop Mask, Global Food-Support Analysis Data (version 0.1) | Multi-satellite | 1000 meters | 2010 | <sup>54</sup> |
| **P10:** Global PALSAR-2/PALSAR Forest/Non-Forest Map (version fnf) | ALOS, ALOS 2 | 25 meters | 2017 | <sup>55</sup> |
| **P11:** Hansen Global Forest Change (version 1.7) | Landsat 8 | 1 arc seconds | 2000 to 2019 | <sup>56</sup> |
| **P12:** Global Forest Canopy Height (version 2005) | Lidar | 30 arc seconds | 2005 | <sup>57</sup> |
| **P13:** JRC Yearly Water Classification History (version 1.2) | Landsat (5,7,8) | 30 meters | 2017 to 2019 | <sup>58</sup> |
| **P14:** JRC Global Surface Water Mapping Layers (version 1.2) | Landsat(5,7,8) | 30 meters | 1984 to 2019 | <sup>58</sup> |
| **P15:** Tsinghua FROM-GLC year of change to impervious surface(version 10) | Landsat | 30 meters | 1985 to 2019 | <sup>59</sup> |

#### 2.1.2 Standardization and Harmonization of LULC legends

Land cover (LC) data describes the main type of natural ecosystem that occupies an area; either by vegetation types such as shrublands, grasslands and forests, or by other biophysical classes such as permanent snow, bare land and water bodies. Land use (LU) includes the way in which people modify or exploit an area, such as in urban areas or agricultural fields.

To build our 29 LULC classes nomenclature, we established a standardization and harmonization approach based on expert knowledge. During this process we took into account the needs of different practitioners in the LULC mapping field and the thematic resolution of the global LULC legends available in GEE. Hence, our nomenclature consists of 23 LC and 6 LU distinct classes interoperable through a set of criteria across 15 LULC products specified in our consensus rules (Table 3). A six-level (L0 to L5) hierarchical structure was adopted in the creation of these 29 LULC classes (Figure 2).

**Table 3.** First stage of the rule set criteria used to find consensus across the 15 Land-Use and Land-Cover (LULC) products available in Google Earth Engine (GEE) for each of the 29 LULC classes contained in the Sentinel2GlobalLULC dataset. P1 to P15: product 1 to 15. C1 to C29: class 1 to class 29. For each product, one or multiple criteria were established to create a global probability map (pixel values 0 or 1) for a given LULC class. A total number of 15×29 = 435 of global probability maps were calculated. The numbers in each column (i.e., from 0 to 220) correspond to the pixel values from each product band. NU: Not Used, NA: Not Available, TC: Tree Cover, G: Tree Gain, L: Tree Loss, D: Datamask, TH: Tree Hight, TCC: Tree Canopy Cover, TCF: Tree-Cover Fraction, and SCF: Shrub-Cover Fraction. ∩:”AND”, ∪:”OR”, +:”ADD”.

|  | P1 | P2 | P3 | P4 | P5 | P6 | P7 | P8 | P9 | P10 | P11 | P12 | P13 | P14 | P15 |
| --- | --- | --- | --- | --- | --- | --- | --- | --- | --- | --- | --- | --- | --- | --- | --- |
| C1 | 16 | 15 | NA | 7 | 11 | 60 | $TCC < 10$ | 200 | 0 | 2 | $(TC < 10) \cap (G = 0) \cap (L = 0) \cap (D \neq 2)$ | $TH < 1$ | 1∪0 | 0 | Not($\geq 1$) |
| C2 | 16 | 15 | NA | 7 | 11 | NA | $TCC < 10$ | 200 ∪ 150 | 0 | 2 | $(TC < 10) \cap (G = 0) \cap (L = 0) \cap (D \neq 2)$ | $TH < 1$ | 1∪0 | 0 | Not($\geq 1$) |
| C3 | 10 | 10 | 1 | 6 | 6 | 30 | $TCC < 10$ | 140 | NA | 2 | $(TC < 10) \cap (G = 0) \cap (L = 0) \cap (D \neq 2)$ | $TH < 2$ | 1∪0 | 0 | Not($\geq 1$) |
| C4 | 7 | 7 | 2 | NA | 5 | $20 \cap (10 < SCF < 50)$ | $TCC < 10$ | 150 | 0 | 2 | $(TC < 10) \cap (G = 0) \cap (L = 0) \cap (D \neq 2)$ | $TH < 2$ | 1∪0 | 0 | Not($\geq 1$) |
| C5 | 6 | 6 | 2 | NA | 5 | $20 \cap (SCF > 50)$ | $TCC < 10$ | 130 | 0 | 2 | $(TC < 10) \cap (G = 0) \cap (L = 0) \cap (D \neq 2)$ | $TH < 2$ | 1∪0 | 0 | Not($\geq 1$) |
| C6 | NA | NA | NA | 4 | 4 | $4 + (15 < TCF < 30)$ | $15 < TCC < 30$ | 60 | NA | 1 | $(15 < TC < 30) \cap (G = 0) \cap (L = 0) \cap (D \neq 2)$ | $TH > 2$ | 1∪0 | 0 | Not($\geq 1$) |
| C7 | NA | NA | NA | 4 | 4 | $4 + (40 < TCF < 60)$ | $40 < TCC < 60$ | 50 | NA | 1 | $(40 < TC < 60) \cap (G = 0) \cap (L = 0) \cap (D \neq 2)$ | $TH > 2$ | 1∪0 | 0 | Not($\geq 1$) |
| C8 | 4 | 4 | 6 | 4 | 4 | $4 + (TCF > 60)$ | $TCC > 60$ | 50 | NA | 1 | $(TC > 60) \cap (G = 0) \cap (L = 0) \cap (D \neq 2)$ | $TH > 2$ | 1∪0 | 0 | Not($\geq 1$) |
| C9 | NA | NA | NA | 3 | 3 | $3 + (15 < TCF < 30)$ | $15 < TCC < 30$ | NA | NA | 1 | $(15 < TC < 30) \cap (G = 0) \cap (L = 0) \cap (D \neq 2)$ | $TH > 2$ | 1∪0 | 0 | Not($\geq 1$) |
| C10 | NA | NA | NA | 3 | 3 | $3 + (40 < TCF < 60)$ | $40 < TCC < 60$ | NA | NA | 1 | $(40 < TC < 60) \cap (G = 0) \cap (L = 0) \cap (D \neq 2)$ | $TH > 2$ | 1∪0 | 0 | Not($\geq 1$) |
| C11 | 3 | 3 | 8 | 3 | 3 | $3 + (TCF > 60)$ | $TCC > 60$ | NA | NA | 1 | $(TC > 60) \cap (G = 0) \cap (L = 0) \cap (D \neq 2)$ | $TH > 2$ | 1∪0 | 0 | Not($\geq 1$) |
| C12 | NA | NA | NA | 2 | 2 | $2 + (15 < TCF < 30)$ | $15 < TCC < 30$ | 40 | NA | 1 | $(15 < TC < 30) \cap (G = 0) \cap (L = 0) \cap (D \neq 2)$ | $TH > 2$ | 1∪0 | 0 | Not($\geq 1$) |
| C13 | NA | NA | NA | 2 | 2 | $2 + (40 < TCF < 60)$ | $40 < TCC < 60$ | 40 | NA | 1 | $(40 < TC < 60) \cap (G = 0) \cap (L = 0) \cap (D \neq 2)$ | $TH > 2$ | 1∪0 | 0 | Not($\geq 1$) |
| C14 | 2 | 2 | 5 | 2 | 2 | $2 + (TCF > 60)$ | $TCC > 60$ | 40 | NA | 1 | $(TC > 60) \cap (G = 0) \cap (L = 0) \cap (D \neq 2)$ | $TH > 2$ | 1∪0 | 0 | Not($\geq 1$) |
| C15 | 9 | 9 | NA | 1 | 1 | $1 + (15 < TCF < 30)$ | $15 < TCC < 30$ | 90 | NA | 1 | $(15 < TC < 30) \cap (G = 0) \cap (L = 0) \cap (D \neq 2)$ | $TH > 2$ | 1∪0 | 0 | Not($\geq 1$) |
| C16 | 8 | 8 | 4 | 1 | 1 | $1 + (40 < TCF < 60)$ | $40 < TCC < 60$ | 70 | NA | 1 | $(40 < TC < 60) \cap (G = 0) \cap (L = 0) \cap (D \neq 2)$ | $TH > 2$ | 1∪0 | 0 | Not($\geq 1$) |
| C17 | 1 | 1 | 7 | 1 | 1 | $1 + (TCF > 60)$ | $TCC > 60$ | 70 | NA | 1 | $(TC > 60) \cap (G = 0) \cap (L = 0) \cap (D \neq 2)$ | $TH > 2$ | 1∪0 | 0 | Not($\geq 1$) |
| C18 | 11 | 11 | NA | NA | NA | 90 | $TCC > 10$ | 170 | NA | NA | $(TC > 10) \cap (G = 0) \cap (L = 0) \cup (D = 2)$ | $TH > 2$ | 2∪3 | 1 | Not($\geq 1$) |
| C19 | 11 | 11 | NA | NA | NA | 90 | $TCC > 10$ | $a.160 \cup 180$<br>$b.Not(170)$ | NA | NA | $(TC > 10) \cap (G = 0) \cap (L = 0) \cup (D = 2)$ | $TH > 2$ | 2∪3 | 1 | Not($\geq 1$) |
| C20 | 11 | 11 | NA | NA | NA | 90 | $TCC < 10$ | $160 \cup 170$<br>$\cup 180$ | NA | NA | $(TC < 10) \cap (G = 0) \cap (L = 0) \cup (D = 2)$ | $TH < 2$ | 2∪3 | 1 | Not($\geq 1$) |
| C21 | 17 | 0 | 0 | 0 | 0 | 200 | NA | 210 | NA | 3 | NA | NA | 3 | 1 | Not($\geq 1$) |
| C22 | 17 | 0 | 0 | 0 | 0 | 80 | NA | 210 | NA | 3 | NA | NA | 3 | 1 | Not($\geq 1$) |
| C23 | 15 | NA | NA | NA | 10 | 70 | NA | 220 | NA | NA | NA | NA | 1∪0 | 0 | Not($\geq 1$) |
| C24 | 12 | 12 | 3∪1 | 5∪6 | 7∪8 | 40 | NA | 11∪14 | $1 \cup 2 \cup 3$<br>$\cup 4 \cup 5$ | NA | NA | NA | 2∪3 | $0 \cup 4 \cup 8 \cup 10$ | Not($\geq 1$) |
| C25 | 12 | 12 | 1 | 6 | 7 | 40 | NA | 11 | $1 \cup 2$ | NA | NA | NA | 1∪0 | 0 | Not($\geq 1$) |
| C26 | 12 | 12 | 1 | 6 | 7 | 40 | NA | 14 | $3 \cup 4 \cup 5$ | NA | NA | NA | 1∪0 | 0 | Not($\geq 1$) |
| C27 | 12 | 12 | 3 | 5 | 8 | 40 | NA | 11 | $1 \cup 2$ | NA | NA | NA | 1∪0 | 0 | Not($\geq 1$) |
| C28 | 12 | 12 | 3 | 5 | 8 | 40 | NA | 14 | $3 \cup 4 \cup 5$ | NA | NA | NA | 1∪0 | 0 | Not($\geq 1$) |
| C29 | 13 | 13 | 10 | 8 | 9 | 50 | NA | 190 | NA | NA | NA | NA | 1∪0 | 0 | NU |

**Figure 2.**
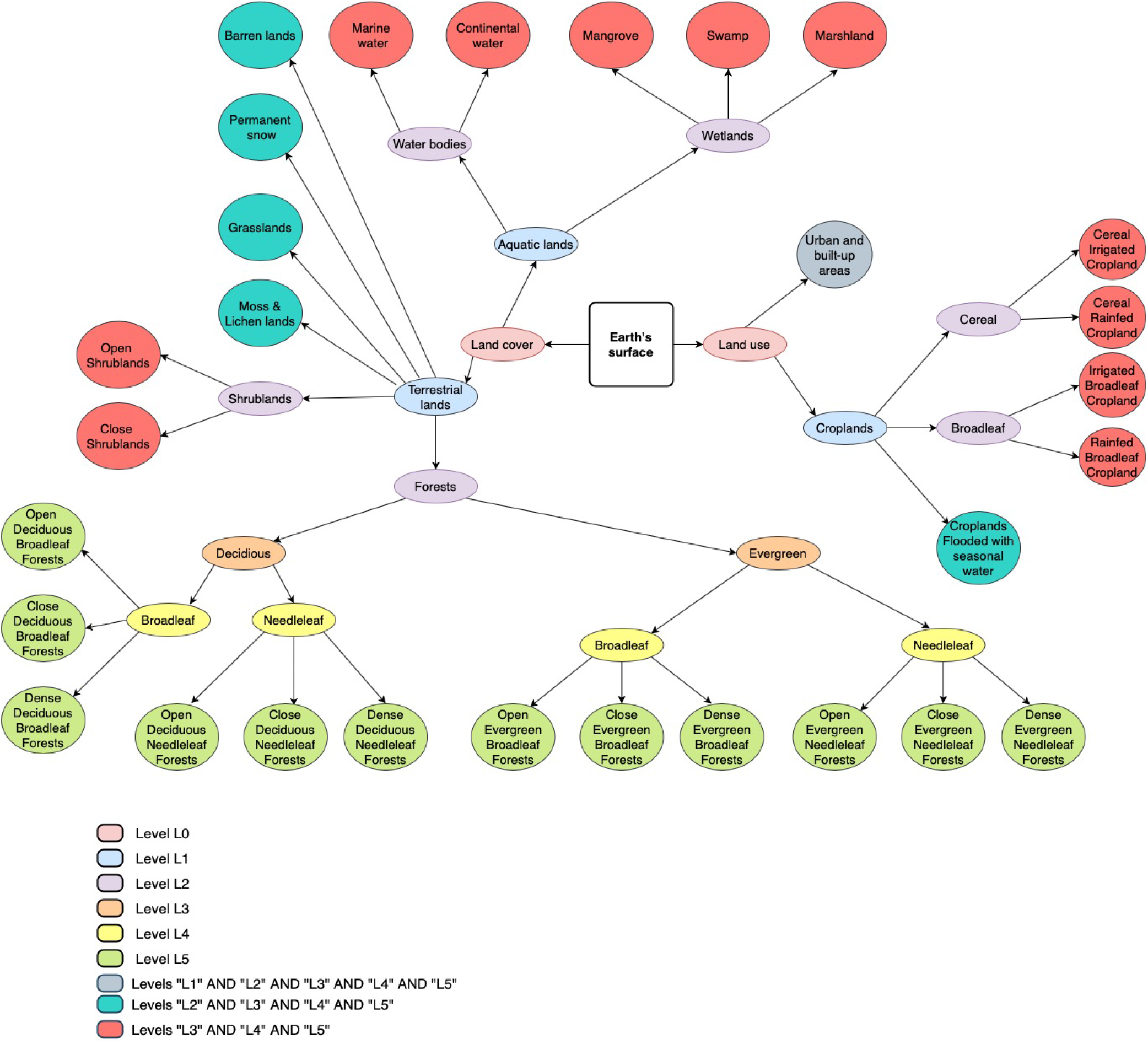
Tree representation of the six-level (L0 to L5) hierarchical structure of the Land-Use and Land-Cover (LULC) classes contained in the Sentinel2GlobalLULC dataset. Outter circular leafs represent the final or most detailed 29 LULC classes of level L5. The followed path to define each class is represented through inner ellipses that contain the names of intermediate classes at different levels between the division of the Earth’s surface (square) into LU and LC (level L0) and the final class circle (level L5). All LULC classes belong to three levels at least, except the 12 forest classes that belong to L5 only.

The LC part contains 20 terrestrial ecosystems and three aquatic ecosystems. The terrestrial systems are: Barren lands, Grasslands, Permanent snow, Moss and Lichen lands, Close Shrublands, Open Shrublands, in addition to 12 Forests classes that differed in their tree cover, phenology, and leaf type. The aquatic classes are: Marine water bodies, Continental water bodies, and Wetlands; furthermore, wetlands are divided into three classes: Marshlands, Mangroves and Swamps. The LU part is composed of urban areas and five coarse cropland types that differed in their irrigation regime and leaf type.

#### 2.1.3 Combining products across time and space

For each one of the 29 LULC classes, we combined in space and time the global LULC information among the 15 GEE LULC products. For each product and LULC type, we first set one or more criteria to create a global mask at the native resolution of the product in which each pixel was classified as 0 or 1 depending on whether it met the criteria for belonging to that LULC type or not, respectively (see first stage in Table 3). Then (see second stage in Table 4), for each LULC type, we calculated the average of all the masks obtained from each product to create a final global probability map at the finest resolution from all products with values ranging between 0 and 1. Value 1 meant that all products agreed to assign that pixel to a particular class and value 0 meant that none of the products assigned it to that particular class (Figure 3). These 0-to-1 values are interpreted as the spatio-temporal purity level of each pixel to belong to a particular LULC class.

**Table 4.** Second stage of the rule set criteria used to find consensus across the 15 Land-Use and Land-Cover (LULC) products available in Google Earth Engine (GEE) for each of the 29 LULC classes contained in the Sentinel2GlobalLULC dataset. P1 to P15: product 1 to 15. C1 to C29: class 1 to class 29. For each LULC class, the 15 global probability maps (with pixel values 0 or 1) obtained in the first stage from products P1 to P15 were spatially combined to build 29 final global probability maps (with pixel values 0 to 1), one for each LULC class (C1 to C29). “Add”:ADD, “*”:MULTIPLY, “Norm”: the normalization using division by number of used products

| Class ID | LULC class | Spatial Combination |
| --- | --- | --- |
| C1 | Barren lands | Norm(Add(P1:P12)*P13*P14*P15) |
| C2 | Moss and Lichen lands | Norm(Add(P1:P12)*P13*P14*P15) |
| C3 | Grasslands | Norm(Add(P1:P12)*P13*P14*P15) |
| C4 | Open Shrublands | Norm(Add(P1:P12)*P13*P14*P15) |
| C5 | Close Shrublands | Norm(Add(P1:P12)*P13*P14*P15) |
| C6 | Open Deciduous Broadleaf Forests | Norm(Add(P1:P12)*P13*P14*P15) |
| C7 | Close Deciduous Broadleaf Forests | Norm(Add(P1:P12)*P13*P14*P15) |
| C8 | Dense Deciduous Broadleaf Forests | Norm(Add(P1:P12)*P13*P14*P15) |
| C9 | Open Deciduous Needleleaf Forests | Norm(Add(P1:P12)*P13*P14*P15) |
| C10 | Close Deciduous Needleleaf Forests | Norm(Add(P1:P12)*P13*P14*P15) |
| C11 | Dense Deciduous Needleleaf Forests | Norm(Add(P1:P12)*P13*P14*P15) |
| C12 | Open Evergreen Broadleaf Forests | Norm(Add(P1:P12)*P13*P14*P15) |
| C13 | Close Evergreen Broadleaf Forests | Norm(Add(P1:P12)*P13*P14*P15) |
| C14 | Dense Evergreen Broadleaf Forests | Norm(Add(P1:P12)*P13*P14*P15) |
| C15 | Open Evergreen Needleleaf Forests | Norm(Add(P1:P12)*P13*P14*P15) |
| C16 | Close Evergreen Needleleaf Forests | Norm(Add(P1:P12)*P13*P14*P15) |
| C17 | Dense Evergreen Needleleaf Forests | Norm(Add(P1:P12)*P13*P14*P15) |
| C18 | Mangrove Wetlands | Norm(Add(P1:P7,P9:P14)*P8*P15) |
| C19 | Swamp Wetlands | Norm(Add(P1:P7,a.P8,P9:P14)*b.P8*P15) |
| C20 | Marshland Wetlands | Norm(Add(P1:P6,P8:P10,P13,P14)*(P11 OR P12 OR P7)*P15) |
| C21 | Marine Water Bodies | Norm(Add(P1:P12)*P13*P14*P15) |
| C22 | Continental Water Bodies | Norm(Add(P1:P12)*P13*P14*P15) |
| C23 | Permanent Snow | Norm(Add(P1:P12)*P13*P14*P15) |
| C24 | Croplands Flooded with seasonal water | Norm(Add(P1:P12)*(P13 OR P14)*P15) |
| C25 | Cereal Irrigated Cropland | Norm(Add(P1:P12)*P13*P14*P15) |
| C26 | Cereal Rainfed Cropland | Norm(Add(P1:P12)*P13*P14*P15) |
| C27 | Irrigated Broadleaf Cropland | Norm(Add(P1:P12)*P13*P14*P15) |
| C28 | Rainfed Broadleaf Cropland | Norm(Add(P1:P12)*P13*P14*P15) |
| C29 | Urban and built-up areas | Norm(Add(P1:P12)*P13*P14*P15) |

**Figure 3.**
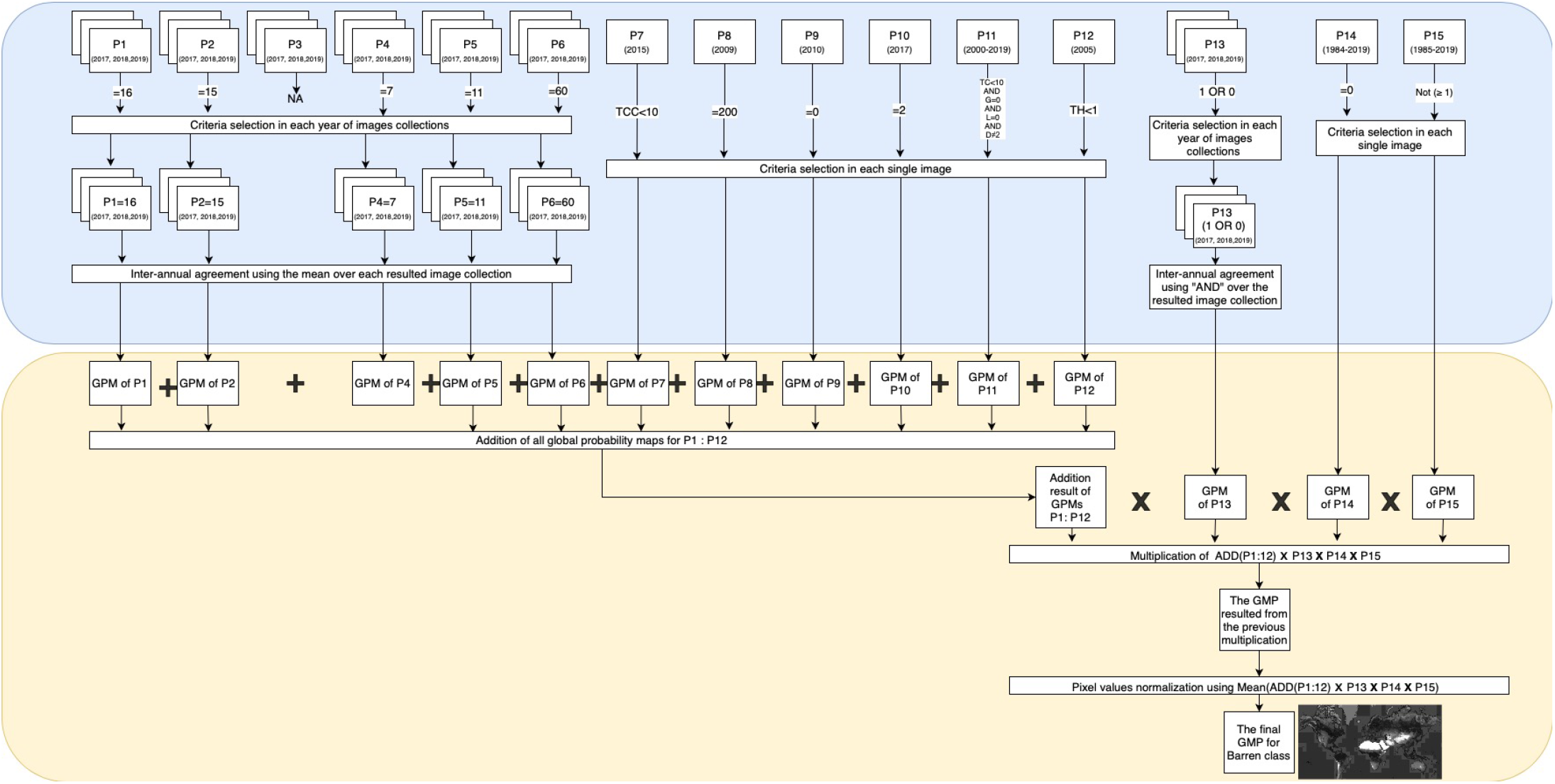
Example of the process of building the final global probability map for one of the 29 Land-Use and Land-Cover (LULC) classes (e.g. C1: “Barren”) by means of spatio-temporal agreement of the 15 LULC products available in Google Earth Engine (GEE). The final map is normalized to values between 0 (white, i.e., areas with no presence of C1 in any product) and 1 (black spots, i.e., areas containing or compatible with the presence of C1 in all 15 products), whereas the shades of grey corresponds to the values in between (i.e., areas that did not contain or were not compatible with the presence of C1 in some of the products). This process is divided into two stages: the first stage (the blue part, see details in Table 3) and the second stage (the yellow part, see details in Table 4). LULC products available for several years are represented with superposed rectangles, while single year products are represented with single rectangles. GMP: global probability map, NA: Not Available.

As an example of the first stage (see details in Table 3), to specify if a given pixel belongs to a dense, evergreen or needleleaf forest, we evaluated its tree cover level using “≤“ and “≥“, while for bands containing the leaf type information, we used the equal operator “=“. For the spatio-temporal combination of multiple criteria we have used the following operators: “AND”,”OR” and “ADD”. For example, we combined the tree cover percentage criteria with the leaf type criteria using “AND” in order to select forest pixels that meets both conditions. To combine many years instances of the same product we used “ADD”, except for product P13 where we used “AND” to select permanent water areas. Whenever we used the “ADD” operator, we normalized pixel values afterwards to bring it back to a probability interval between 0 and 1 using the division by the total number of combined years or criteria.

In the second stage (see details in Table 4), we combined for each LULC class, the 15 global probability maps resulted from the previous stage to create a final global probability map. This combination was carried out using various operators such as “ADD”, “MULTIPLY” and “OR”, depending on the LULC type. When “ADD” was used, the final pixel values were normalized by dividing the final addition value of each pixel by the total number of added products. The “MULTIPLY” operator was mostly used at the end, to remove urban areas from non-urban LULC classes, or to remove water from non-water systems. The multiplication operator was also adopted to make sure that a certain criteria was respected in the final probability map. For instance, for the swamp class, we multiplied all pixels in the final stage by a water mask where saline water areas have a value of 0 in order to eliminate mangroves from swamp pixels and vice versa. Finally, we used “OR” operator between different water related products in order to take advantage of the fact that each one complements the other in terms of spatial coverage and accuracy.

#### 2.1.4 Re-projection and Selection of purity threshold

After the consensus assessment, the 29 final probability maps maintained the spatial resolution of the last aggregated LULC product, i.e., the water product at 30m/pixel. Since our objective was finding pure tiles of 224 × 224 10-m pixels (i.e. Sentinel-2 pixels) for each LULC class, we reprojected the 30 m/pixel probability maps to 2240 m/pixel by using the spatial mean reducer in GEE.

For each one of the reprojected maps, we defined a pixel value threshold to decide whether a given 2240 × 2240 m tile was representative of each LULC class or not. If the number of available pure tiles (i.e., pixel value = 1) was too small for one class, we decreased the threshold for purity level for that class until getting a large enough number of tiles (the purity level is always provided as metadata for each tile). On the other hand, when the number of pure tiles for a LULC class was too large, (i.e., greater than 14000), we applied a stratified selection to download a maximum of 14000 images. This selection was carried out through an automatic maximum geographic distance algorithm to guarantee that selected images were as geographically far from each other as possible. In Table 5, we present the number of tiles we found and downloaded for each LULC class using thresholds ranging from 0.75 to 1. We illustrated the reprojection and selection processes in Figure 4.

**Table 5.** Summary of the varying number of found and eventually selected Sentinel-2 image tiles of 224 × 224 pixels depending on the different consensus level reached across the 15 Land-Use and Land-Cover (LULC) products available in Google Earth Engine (GEE) for each of the 29 LULC classes contained in the Sentinel2GlobalLULC dataset. LULC classes that due to the too large number of samples had to undergo a stratified selection by maximizing geographical distance among samples are highlighted in bold.

| LCLU Class | Consensus probability values (%) |  |  |  |  |  | Number of selected images | Stratified selection |
| --- | --- | --- | --- | --- | --- | --- | --- | --- |
|  | 0.75 (75%) | 0.80 (80%) | 0.85 (85%) | 0.90 (90%) | 0.95 (95%) | 1.00 (100%) |  |  |
| Urban | 63953 | - | 34102 | 21814 | 12590 | 192 | 12590 | no |
| Barren | 4330418 | - | 4055836 | 3876467 | 3545756 | 2668009 | **14000 (2668009)** | yes |
| Moss and Lichen | 59120 | - | 18438 | 4669 | 1158 | 0 | 4669 | no |
| Close Shrublands | 41407 | 12502 | 1872 | 226 | 16 | 0 | 12502 | no |
| Open Shrublands | 2461415 | - | 1209375 | 644272 | 101288 | 805 | **14000 (101288)** | yes |
| Marshland | 4205 | - | 675 | 143 | 15 | 0 | 4205 | no |
| Swamp | 489 | - | 4 | 0 | 0 | 0 | 489 | no |
| Mangrove | 425 | - | 63 | 3 | 0 | 0 | 425 | no |
| Grassland | 4022949 | - | 1894337 | 943177 | 128263 | 8895 | 8895 | no |
| Rainfed Broadleaf Cropland | 427314 | - | 209143 | 99337 | 32123 | 416 | 416 | no |
| Irrigated Broadleaf Cropland | 224867 | - | 92488 | 53064 | 30691 | 354 | 354 | no |
| Cereal Rainfed Cropland | 1185497 | - | 604459 | 284914 | 91147 | 1022 | 1022 | no |
| Cereal Irrigated Cropland | 517789 | - | 167994 | 52959 | 23555 | 842 | 842 | no |
| Cropland Seasonal Water | 6048 | - | 3192 | 2008 | 995 | 15 | 2008 | no |
| Dense Evergreen Needleleaf Forest | 474138 | - | 178293 | 66151 | 13995 | 0 | 13995 | no |
| Close Evergreen Needleleaf Forest | 43040 | 3875 | 69 | 0 | 0 | 0 | 3875 | no |
| Open Evergreen Needleleaf Forest | 17462 | 3939 | 331 | 0 | 0 | 0 | 3939 | no |
| Dense Evergreen Broadleaf Forest | 2131269 | - | 1829897 | 1594657 | 1232914 | 144026 | **14000 (144026)** | yes |
| Close Evergreen Broadleaf Forest | 12512 | 1270 | 77 | 1 | 0 | 0 | 1270 | no |
| Open Evergreen Broadleaf Forest | 574 | 42 | 0 | 0 | 0 | 0 | 574 | no |
| Dense Deciduous Needleleaf Forest | 60866 | - | 12954 | 2888 | 148 | 0 | 2888 | no |
| Close Deciduous Needleleaf Forest | 42166 | 6383 | 35 | 0 | 0 | 0 | 6383 | no |
| Open Deciduous Needleleaf Forest | 10439 | 23 | 0 | 0 | 0 | 0 | 10439 | no |
| Dense Deciduous Broadleaf Forest | 399264 | - | 176176 | 97182 | 31284 | 1 | **14000 (31284)** | yes |
| Close Deciduous Broadleaf Forest | 71127 | - | 1353 | 23 | 1 | 0 | 1353 | no |
| Open Deciduous Broadleaf Forest | 25342 | 4439 | 466 | 2 | 0 | 0 | 4439 | no |
| Permanent Snow | 1065127 | - | 1033466 | 1013490 | 984014 | 877232 | **14000 (877232)** | yes |
| Continental Water Bodies | 3543953 | - | 3199652 | 343779 | 318483 | 265214 | **14000 (265214)** | yes |
| Marine Water Bodies | 3606955 | - | 3357810 | 2903459 | 2822544 | 2577444 | **14000 (2577444)** | yes |

**Figure 4.**
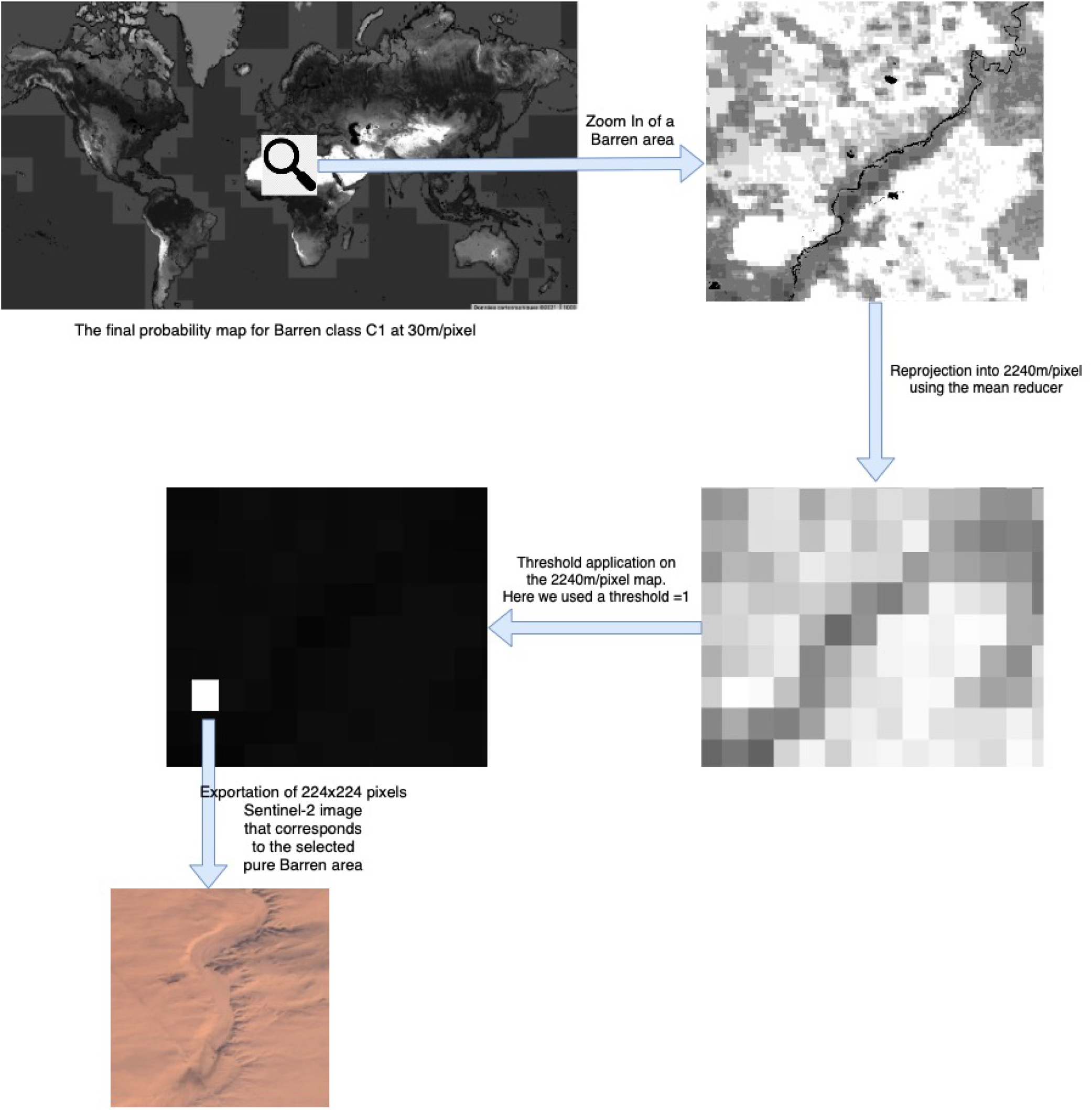
Example of the workflow to obtain a Sentinel-2 image tile of 2240 × 2240 m for one of the 29 Land-Use and Land-Cover (LULC) classes (e.g. C1: “Barren”). The process starts with the reprojected final global probability map obtained from stage two (Table 4) and ends with its exportation to the repository of a Sentinel-2 image tile of 224 × 224 pixels. The white rectangle is the only one having a probability value of 1 (Recall that the purity threshold used for Barren was 1, i.e., 100%). The black pixels has a null probability value, while the probability values between 0 and 1 are represented in gray scale levels.

### 2.2 Data Extraction

Sentinel2GlobalLULC provides the user with two types of data: CSV files and Sentinel-2 RGB images. In the following subsections, we first present the additional gHM index attached to these both data types, then the adopted methods to generate each one of them.

#### 2.2.1 gHM values extraction

As an additional metadata related to the level of human influence in each image, we calculated for each tile the spatial mean of the global human modification index for terrestrial lands^41^, where 0 means no human modification and 1 means complete transformation. Since the original gHM product was mapped at 1 × 1 km resolution, we reprojected it to 2240 × 2240 m using the same procedure than explained for the LULC consensus masks.

#### 2.2.2 CSV files generation

Once we identified tiles to be selected for each LULC class, we have grouped their center coordinates into a CSV file each. Tiles were organized giving their probability values in an descendant order. Each row in the CSV file corresponds to a selected tile in that class. In fact, these CSV files contains the geographical center point coordinates, the pixel purity value, the name of the attributed LULC class in addition to the extracted gHM value for that tile. Then, we used the geographical coordinates of each tile to identify its exact administrative address geolocation. To implement this reverse geo-referencing operation, we used a free request-unlimited python module called [reverse_geocoder](https://pypi.org/project/reverse-geocode/). This method has allowed us to identify the country code, the administrative department at two levels and the locality of each tile in the CSV files. This way, we integrated in all LULC classes CSV files these reverse geo-referencing information as new columns.

For LULC class that has more than 14000 pure tiles, we have released the coordinates before and after the stratified selection in case the user was interested in all tiles and not only the exported ones. These coordinates could allow the end user to download new images if needed.

#### 2.2.3 Sentinel-2 RGB images exportation

After extracting all these pieces of information and grouping them into CSV files, we went back to the geographic center coordinates of each tile and used them to extract the corresponding 224 × 224 pixel Sentinel-2 RGB tiles using GEE. Each exported image was identical to the 2240 × 2240 m area covered by its Sentinel-2 tile.

We chose “Sentinel-2 MSI (Multi-Spectral Instrument) product” since it is free and publicly available in GEE at the fine resolution of 10 × 10 m. We chose “Level-1C” since it provides the longest data availability of Sentinel-2 images. To build RGB images, we extracted the three bands B4, B3 and B2 that correspond to Red, Green and Blue channels, respectively.

To minimize the effect of atmospheric effects on the RGB images, such as clouds, aerosols, smoke, etc., every image was built from the 25th-percentile aggregation of its corresponding image collection gathered by Sentinel-2 satellites between June 2015 and October 2020. In addition, we previously discarded all pixels where the maximum cloud probability exceeded 20% according to the metadata provided in the Sentinel-2 collection.

Usually, Sentinel-2 MSI product includes true colour images in JPEG2000 format, except for the “Level-1C” collection used here. The three original bands (B4, B3, and B2) required a saturation stretching of their reflectance values into 0-255 RGB digital values. Thus, we stretched the saturation reflectance of 3558 into 255 to obtaine true RGB channels with digital values between 0 and 255. The choice of these mapping numbers was taken from the Sentinel-2 true colour image recommendations section of [Sentinel user guidelines](https://sentinel.esa.int/web/sentinel/user-guides/sentinel-2-msi/definitions). Finally, after exporting the selected tiles for each LULC class as “.tif” images, we converted them into “.jpeg” format using a lossless conversion algorithm.

### 2.3 Technical implementation

To implement all our methodology steps, we first created a javascript in GEE for each LULC class. Each script is a multi-task javascript where we implemented a switch command to control which task we want to execute. In each one of these scripts, we selected from [GEE LULC datasets repository](https://developers.google.com/earth-engine/datasets/) the 15 LULC products used to build the consensus of that LULC class. Each script was responsible of elaborating the spatio-temporal combination of the selected products and generating the final consensus map for that LULC class as described in the subsection Combining products across time and space. Then, it exports the final global probability map as an asset into GEE server storage to make its reprojection faster. In the same script, once the consensus map exportation was done, we imported it from the GEE assets storage and reprojected it to 2240 × 2240 m resolution; then, we exported the new reprojected map into GEE assets storage again to make its analysis and processing faster. Afterwards, we imported the reprojected map into the same script and apply different processing tasks. During this processing phase, many purity threshold values were evaluated. Then, we elaborated in this same script the pure tiles identification and their center coordinates exportation into a CSV file. A distinct GEE script was developed to import, reproject and export the global gHM map. The resulted gHM map was saved as an asset, then imported and used in each one of the 29 LULC multi-task scripts.

A python script was developed separately to read the exported CSV files for each LULC class and apply the reverse geo-referencing on their pure tiles coordinates then add the found geolocalization data (country code, locality…etc) to the original CSV files as new columns. Then, another python script was implemented to read the new resulted CSV files with all their added columns (reverse geo-referencing data, gHM data) and use the center coordinates of each pure tile in that class to export its corresponding Sentinel-2 satellite tif image within GEE through the python API. Finally, after downloading all the exported tif images from our Google drive, we created another python script to convert the exported tif images into JPEG format.

#### Data Records

Sentinel2GlobalLULC dataset is stored in the following Zenodo repository(DOI:10.5281/zenodo.5055632). This dataset consists of three zip compressed folders:

- Sentinel-2 GeoTiff images folder: This folder contains the exported Sentinel-2 RGB images for each LULC class grouped into sub-folders named according to each LULC class. Each image has a filename with the following structure: “LULC class ID_LULC class short name_Pixel probability value_Image ID_GHM value_Latitude _Longitude_Country code_Administrative department level1_Administrative department level2_Locality”. Pixel probability value can be interpreted as the spatial purity of the image to represent that LULC class and was calculated as the spatial mean of all the pixels of the final probability maps contained in each image tile, reprojected and expressed as a percentage. Short names for all classes were derived from the original ones in a way to have exactly 13 characters each, and IDs for different classes were assigned randomly. This information for each class is explained in Table 6.
- Sentinel-2 JPEG images folder: This folder contains the same images as in the GeoTiff folder, but converted into “.jpeg” format while preserving the same nomenclature and organization. In Figure 5, we illustrate a sample of each one of the 29 classes images in JPEG format.
- CSV files folder: For user convenience, the metadata of every image tile (i.e., the same information already contained in the image filenames) is also provided in CSV format. Image tiles in the CSV files are organized from the highest to the lowest consensus probability value. These CSV files have 11 columns: ID of LULC Class, Short name of LULC Class, ID Image, Pixel Probability Value as percentage, GHM Value, Center Latitude, Center Longitude, Country Code, Administative Departement Level 1, Administative Departement Level 2, Locality.

For too large LULC classes (i.e., with more than 14000 potential samples) that had to undergo a stratified selection, we provide the user with 2 CSV files: one containing all pure tiles coordinates without geo-referencing columns, and another file just containing the 14000 exported tiles coordinates with their geo-referencing information.

**Table 6.** Dictionary to map each Land-Use and Land-Cover (LULC) class to its corresponding short name and ID in the Sentinel2GlobalLULC dataset

| LCLU Class | Short name | Class ID |
| --- | --- | --- |
| Urban | UrbanBIUpArea | 29 |
| Barren | BarrenLands__ | 1 |
| Moss and Lichen | MossAndLichen | 2 |
| Close Shrublands | SrublandClose | 5 |
| Open Shrublands | ShrublandOpen | 4 |
| Marshland | WetlandMarshl | 20 |
| Swamp | WetlandSwamps | 19 |
| Mangrove | WetlandMangro | 18 |
| Grassland | Grasslands__ | 3 |
| Rainfed Broadleaf Cropland | CropBroadRain | 28 |
| Irrigated Broadleaf Cropland | CropBroadIrri | 27 |
| Cereal Rainfed Cropland | CropCereaRain | 26 |
| Cereal Irrigated Cropland | CropCereaIrri | 25 |
| Cropland Seasonal Water | CropSeasWater | 24 |
| Dense Evergreen Needleleaf Forest | ForestsDeEvNe | 17 |
| Close Evergreen Needleleaf Forest | ForestsClEvNe | 16 |
| Open Evergreen Needleleaf Forest | ForestsOpEvNe | 15 |
| Dense Evergreen Broadleaf Forest | ForestsDeEvBr | 14 |
| Close Evergreen Broadleaf Forest | ForestsClEvBr | 13 |
| Open Evergreen Broadleaf Forest | ForestsOpEvBr | 12 |
| Dense Deciduous Needleleaf Forest | ForestsDeDeNe | 11 |
| Close Deciduous Needleleaf Forest | ForestsClDeNe | 10 |
| Open Deciduous Needleleaf Forest | ForestsOpDeNe | 9 |
| Dense Deciduous Broadleaf Forest | ForestsDeDeBr | 8 |
| Close Deciduous Broadleaf Forest | ForestsClDeBr | 7 |
| Open Deciduous Broadleaf Forest | ForestsOpDeBr | 6 |
| Permanent Snow | PermanentSnow | 23 |
| Continental Water Bodies | WaterBodyCont | 22 |
| Marine Water Bodies | WaterBodyMari | 21 |

**Figure 5.**
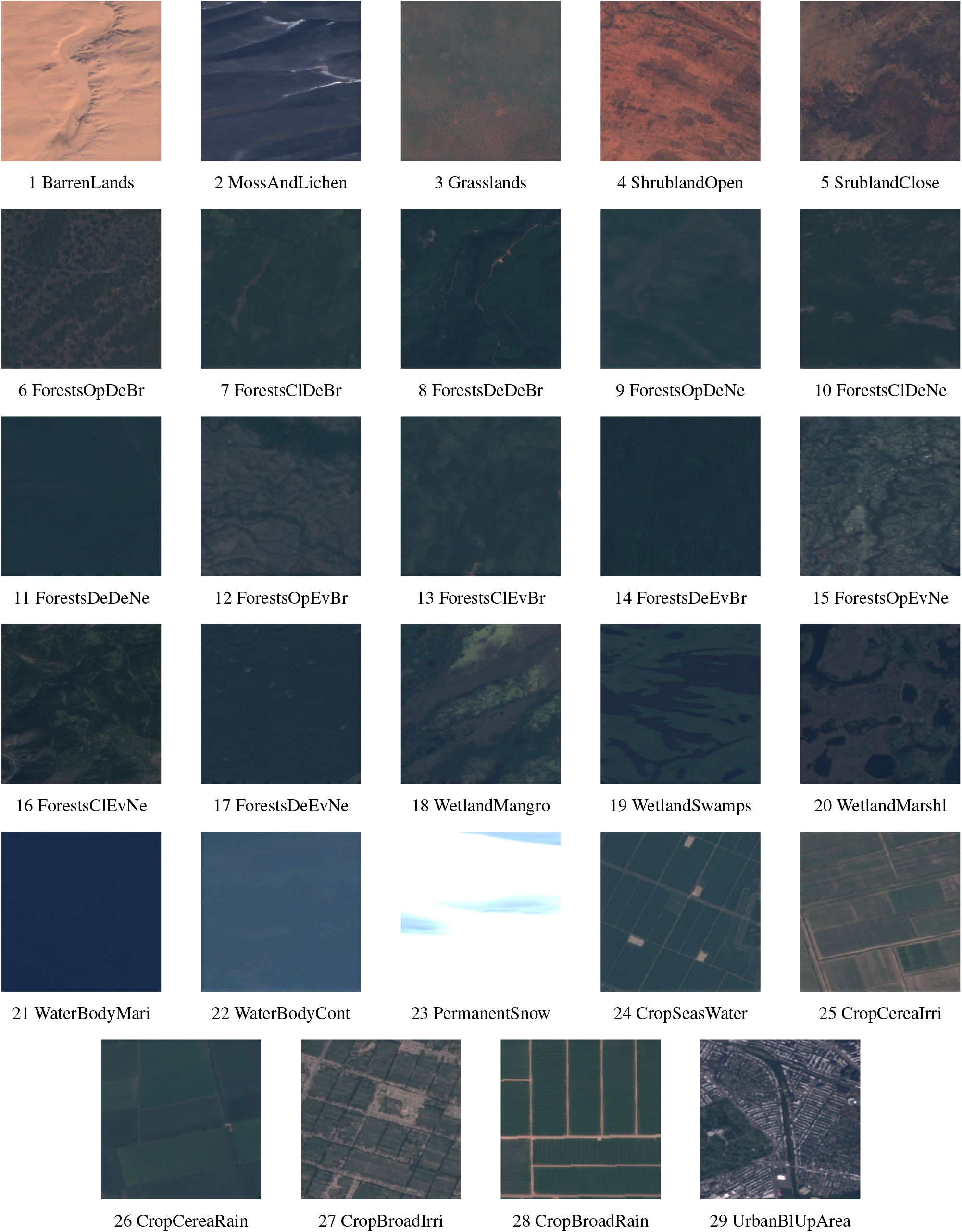
Samples of images for each one of the 29 Land-Use and Land-Cover (LULC) classes contained in the Sentinel2GlobalLULC dataset

#### Technical Validation

To assess the quality of the Sentinel2GlobalLULC dataset in terms of its representativity of each LULC class and of image quality, two of the coauthors visually inspected very high resolution imagery (Google Earth and Bing Maps) of a random sample of each class. The validation process was established in three stages:

- First, for each LULC class, we selected 100 samples to visually verify their LULC annotation. To maximize the global representativity of the validated samples, the selection of these 100 samples was carried out by maximizing the geographical distance among all samples using an add-hoc script in R. In Figure7, we present the distribution map of the 100 samples selected for each LULC class.
- Second, each one of the selected samples was visually inspected in Google Earth and Bing Maps by two of the co-authors (E.G. and D.A-S.) to independently assign it to one of the 29 LULC classes. These two experts assigned each sample to a LULC class when it occupied more than 70% of the image tile.
- Third, the confusion matrix for this validation was calculated at six different levels of our LULC classification hierarchy (from L0 to L5 as presented in Figure2). In Table7, we summarized the obtained F1 scores at each level.

**Figure 6.**
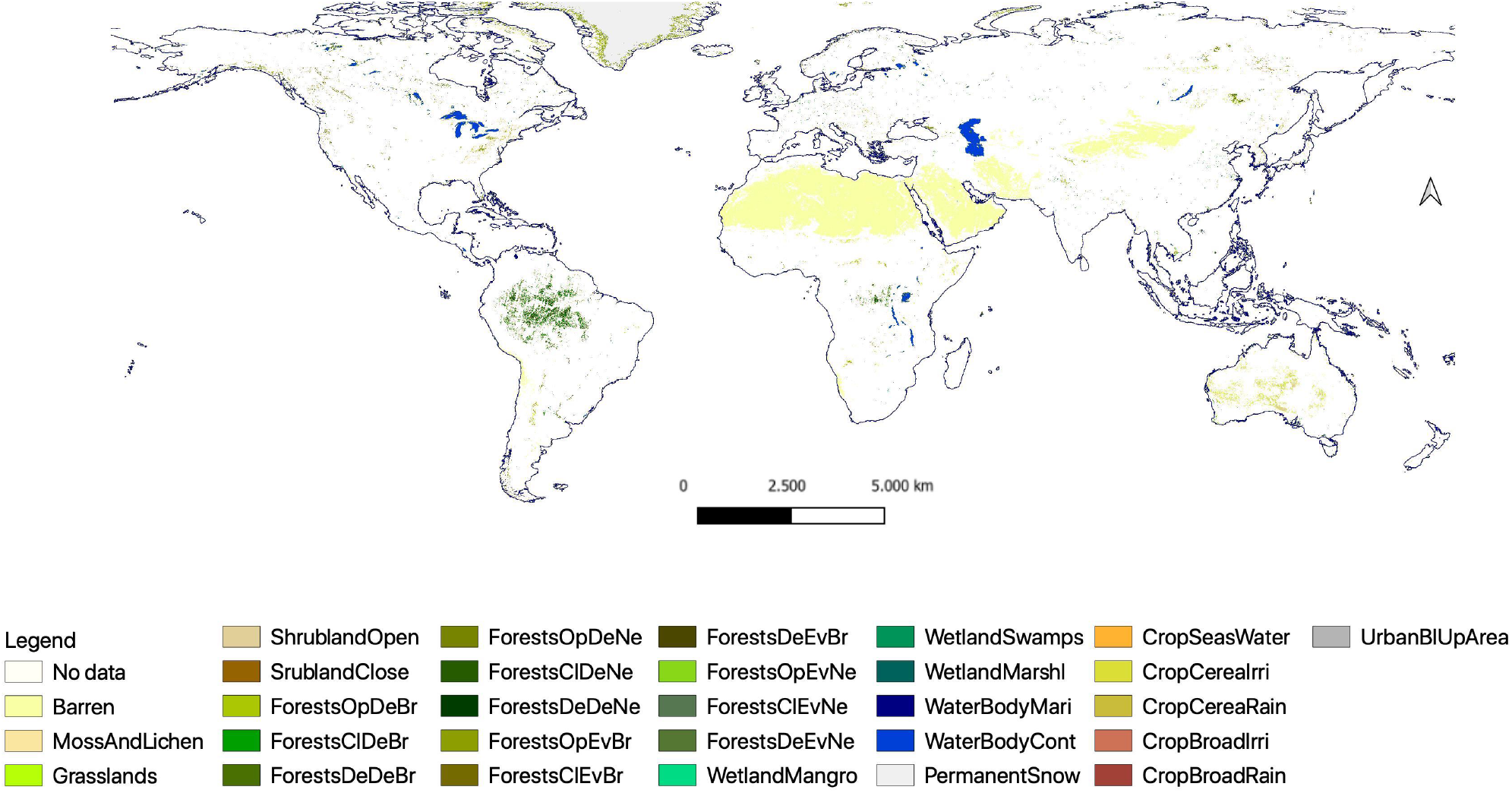
Global map of the distribution of the 2240 × 2240 m tiles representing 29 Land-Use and Land-Cover (LULC) classes that were generated from the spatio-temporal agreement across the 15 global LULC products available in Google Earth Engine. The purity threshold used for each LULC class is specified in Table 5.

**Figure 7.**
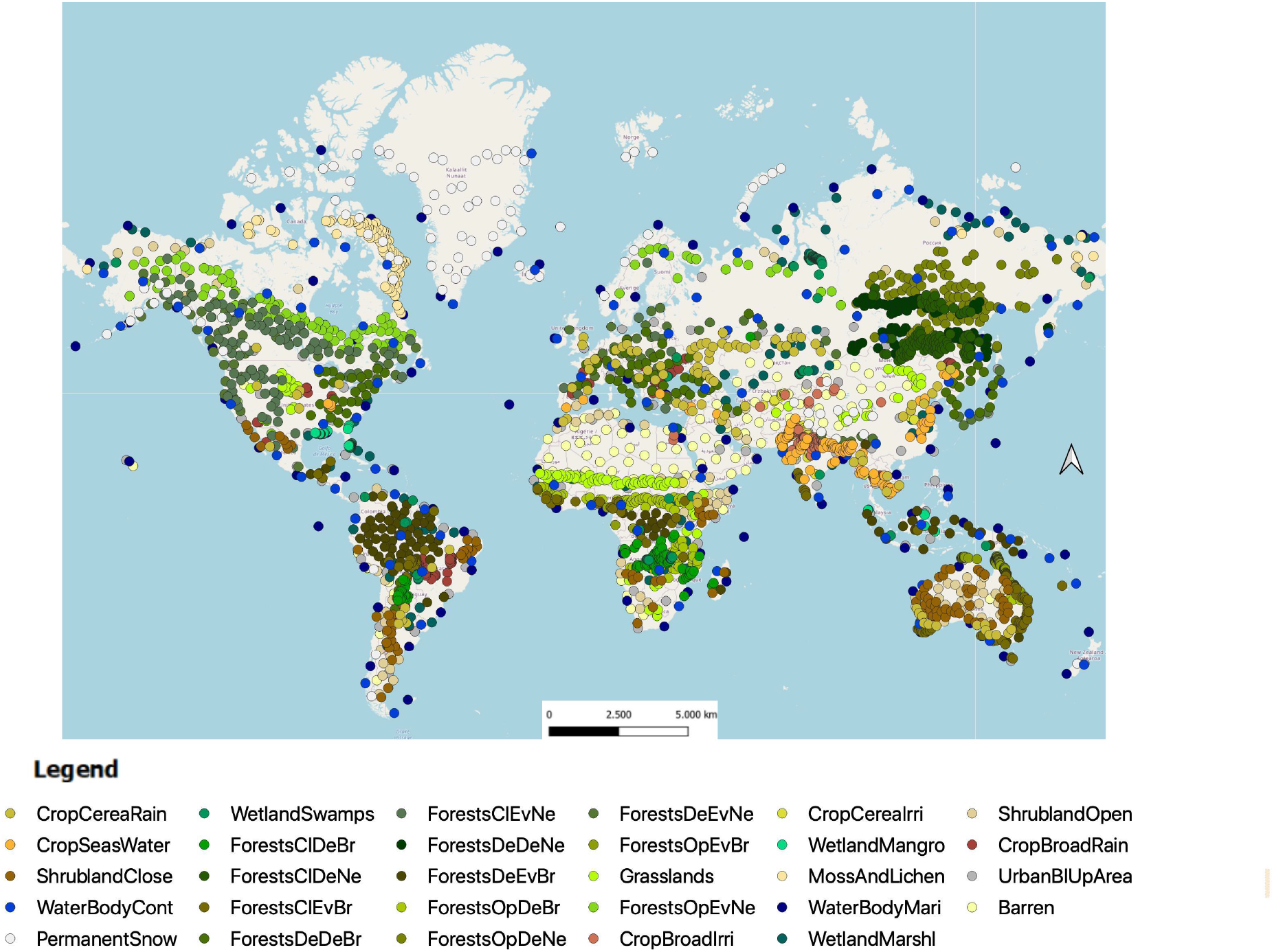
Global distribution of the selected 100 images for each Land-Use and Land-Cover (LULC) class to perform the validation of the 29 LULC classes contained in the Sentinel2GlobalLULC dataset. An add-hoc script in R was used to maximize the geographical distance among the 100 points of each class.

**Table 7.** Results of the validation procedure of the representativeness of the images contained in the Sentinel2GlobalLULC dataset for each Land-Use and Land-Cover (LULC) class at different levels of the hierarchical legend (from L0 to L5). Accuracy is expressed as the mean F1 score (i.e., a balance between precision and recall) for each LULC class at each level, rounded to two decimal values.

| L0 | F1 | L1 | F1 | L2 | F1 | L3 | F1 | L4 | F1 | L5 | F1 |
| --- | --- | --- | --- | --- | --- | --- | --- | --- | --- | --- | --- |
| Land Cover | 0.99 | Terrestrial Lands | 1.00 | BarrenLands | 0.97 | BarrenLands | 0.97 | BarrenLands | 0.97 | BarrenLands | 0.97 |
|  |  |  |  | MossAndLichen | NA | MossAndLichen | NA | MossAndLichen | NA | MossAndLichen | NA |
|  |  |  |  | Grasslands | 0.75 | Grasslands | 0.75 | Grasslands | 0.75 | Grasslands | 0.75 |
|  |  |  |  | Shrubland | 0.89 | ShrublandOpen | 0.76 | ShrublandOpen | 0.76 | ShrublandOpen | 0.76 |
|  |  |  |  |  |  | SrublandClose | 0.97 | SrublandClose | 0.97 | SrublandClose | 0.97 |
|  |  |  |  | Forests | 1.00 | ForestsDe | 1.00 | ForestsDeBr | 1.00 | ForestsOpDeBr | 0.82 |
|  |  |  |  |  |  |  |  |  |  | ForestsCIDEBr | 0.89 |
|  |  |  |  |  |  |  |  |  |  | ForestsDeDeBr | 0.96 |
|  |  |  |  |  |  |  |  | ForestsDeNe | 1.00 | ForestsOpDeNe | 0.92 |
|  |  |  |  |  |  |  |  |  |  | ForestsCIDEne | 0.88 |
|  |  |  |  |  |  |  |  |  |  | ForestsDeDeNe | 0.95 |
|  |  |  |  |  |  | ForestsEv | 0.99 | ForestsEvBr | 0.99 | ForestsOpEvBr | 0.70 |
|  |  |  |  |  |  |  |  |  |  | ForestsCIEvBr | 0.72 |
|  |  |  |  |  |  |  |  |  |  | ForestsDeEvBr | 0.91 |
|  |  |  |  |  |  |  |  | ForestsEvNe | 1.00 | ForestsOpEvNe | 0.82 |
|  |  |  |  |  |  |  |  |  |  | ForestsCIEvNe | 0.88 |
|  |  |  |  |  |  |  |  |  |  | ForestsDeEvNe | 0.99 |
|  |  |  |  | PermanentSnow | 1.00 | PermanentSnow | 1.00 | PermanentSnow | 1.00 | PermanentSnow | 1.00 |
|  |  | Aquatic Lands | 0.98 | Wetland | 0.96 | WetlandMangro | 0.96 | WetlandMangro | 0.96 | WetlandMangro | 0.96 |
|  |  |  |  |  |  | WetlandSwamps | 0.99 | WetlandSwamps | 0.99 | WetlandSwamps | 0.99 |
|  |  |  |  |  |  | WetlandMarshl | 0.94 | WetlandMarshl | 0.94 | WetlandMarshl | 0.94 |
|  |  |  |  | WaterBody | 0.99 | WaterBodyMari | 0.95 | WaterBodyMari | 0.95 | WaterBodyMari | 0.95 |
|  |  |  |  |  |  | WaterBodyCont | 0.93 | WaterBodyCont | 0.93 | WaterBodyCont | 0.93 |
| Land Use | 0.98 | Croplands | 0.98 | CropSeasWater | 0.93 | CropSeasWater | 0.93 | CropSeasWater | 0.93 | CropSeasWater | 0.93 |
|  |  |  |  | CropCerea | 0.99 | CropCereaIrri | 1.00 | CropCereaIrri | 1.00 | CropCereaIrri | 1.00 |
|  |  |  |  |  |  | CropCereaRain | 0.98 | CropCereaRain | 0.98 | CropCereaRain | 0.98 |
|  |  |  |  | CropBroad | 0.99 | CropBroadIrri | 1.00 | CropBroadIrri | 1.00 | CropBroadIrri | 1.00 |
|  |  |  |  |  |  | CropBroadRain | 0.99 | CropBroadRain | 0.99 | CropBroadRain | 0.99 |
|  |  | UrbanBIUpArea | 0.99 | UrbanBIUpArea | 0.99 | UrbanBIUpArea | 0.99 | UrbanBIUpArea | 0.99 | UrbanBIUpArea | 0.99 |
| Mean | 0.99 |  | 0.98 |  | 0.95 |  | 0.95 |  | 0.95 |  | 0.91 |

The obtained mean F1 scores ranged from 0.99 at level L0 to 0.91 at level L5 (Table7). Such decrease in accuracy as the number of classes increased from level L0 to level L5 was mainly due to the hard distinction between forest types at L5 and the complexity of visual features in Grasslands and Shrublands classes from level L2.

#### Usage Notes

To make the Sentinel2GlobalLULC dataset easier to use, reproduce, and exploit and to promote its usage with DL models, we have provided users with a python code to load all RGB images and train several Convolutional Neural Networks (CNNs) models on them using different learning hyper-parameters. Knowing that most CNN frameworks admit only jpeg or png images formats, we provided a python script to convert “.tif” into “.jpeg” format with a full control on the conversion quality and the choice of images to convert. Moreover, as for some LULC classes we limited the number of exported images to 14000, we have provided a python script that can help the user to export more Sentinel-2 images of these classes if needed, using the coordinates stored in the CSV files.

In addition, to provide a global insight about the consistency and accuracy of the global distribution of these 29 LULC classes, we also publicly shared the final reprojected global consensus maps for each class as GEE assets. To help the user to visualize the global distribution of each LULC class, we have provided a GEE script with the assets links to choose, import, manipulate, and visualize any LULC class map. Image exportation is also possible through python API for GEE and we gave the user a complete control on the number of tiles to export, the time interval to select for image collections, the cloud removal parameters, the true RGB colors calibration, and the Google drive account where to store the exported images. The user should be aware that GEE imposes a limited request number with a maximum of 3000 exportation tasks to run simultaneously on the same Google account.

## Code Availability

All used scripts to implement our dataset and links to the GEE stored assets are available in the following Github repository (DOI:10.5281/zenodo.5638409) with guidelines stored in a README file explaining all instructions about their execution.

## Acknowledgements

This work was supported by projects A-RNM-256-UGR18 and A-TIC-458-UGR18A, and is part of LifeWatch SmartEco-Mountains project, all partially funded by the European Union Funds for Regional Development. S.T. was supported by the Ramón y Cajal Programme (No. RYC-2015-18136). S.T., D.A.-S., and F.H. were supported by the project DeepSCOP-Ayudas Fundación BBVA a Equipos de Investigación Científica en Big Data 2018. E.G. was supported by the European Research Council grant agreement n° 647038 (BIODESERT).

## Author contributions statement

Y.B. contributed to the conception of the dataset, implemented the code, performed all the data extraction and wrote the paper. D.A.-S. contributed to the conception and validation of the dataset, provided guidance, and wrote the paper. R.K. contributed to the conception of the dataset. E.G. validated the dataset. F.H. and B.A. provided edits and suggestions. S.T. contributed to the conception of the dataset and wrote the manuscript.

## References

1. Di Gregorio, A. Land cover classification system: classification concepts and user manual: LCCS, vol. 2 (Food & Agriculture Org., 2005).

2. Pielke, R. A. et al. Interactions between the atmosphere and terrestrial ecosystems: influence on weather and climate. Global change biology 4, 461–475 (1998).

3. Menke, S., Holway, D., Fisher, R. & Jetz, W. Characterizing and predicting species distributions across environments and scales: Argentine ant occurrences in the eye of the beholder. Global Ecology Biogeography 18, 50–63 (2009).

4. Verburg, P. H., Neumann, K. & Nol, L. Challenges in using land use and land cover data for global change studies. Global change biology 17, 974–989 (2011).

5. DeFries, R. Terrestrial vegetation in the coupled human-earth system: contributions of remote sensing. Annual Review Environment Resources 33, 369–390 (2008).

6. Pfeifer, M., Disney, M., Quaife, T. & Marchant, R. Terrestrial ecosystems from space: a review of earth observation products for macroecology applications. Global Ecology Biogeography 21, 603–624 (2012).

7. Quaife, T. et al. Impact of land cover uncertainties on estimates of biospheric carbon fluxes. Global Biogeochemical Cycles 22 (2008).

8. Herold, M. et al. A joint initiative for harmonization and validation of land cover datasets. IEEE Transactions on Geoscience Remote Sensing 44, 1719–1727 (2006).

9. Townshend, J., Justice, C., Li, W., Gurney, C. & McManus, J. Global land cover classification by remote sensing: present capabilities and future possibilities. Remote Sensing Environment 35, 243–255 (1991).

10. Loveland, T. R. et al. Development of a global land cover characteristics database and igbp discover from 1 km avhrr data. International Journal Remote Sensing 21, 1303–1330 (2000).

11. Bartholome, E. & Belward, A. S. Glc2000: a new approach to global land cover mapping from earth observation data. International Journal Remote Sensing 26, 1959–1977 (2005).

12. Tuanmu, M.-N. & Jetz, W. A global 1-km consensus land-cover product for biodiversity and ecosystem modelling. Global Ecology Biogeography 23, 1031–1045 (2014).

13. Sheng, G., Yang, W., Xu, T. & Sun, H. High-resolution satellite scene classification using a sparse coding based multiple feature combination. International journal remote sensing 33, 2395–2412 (2012).

14. Xia, G. et al. Aid: A benchmark dataset for performance evaluation of aerial scene classification. arxiv 2016. arXiv preprint 1608.05167.

15. Xia, G.-S. et al. Structural high-resolution satellite image indexing. In ISPRS TC VII Symposium-100 Years ISPRS, vol. 38, 298–303 (2010).

16. Zhao, L., Tang, P. & Huo, L. Feature significance-based multibag-of-visual-words model for remote sensing image scene classification. Journal Applied Remote Sensing 10, 035004 (2016).

17. Zhou, W., Newsam, S., Li, C. & Shao, Z. Patternnet: A benchmark dataset for performance evaluation of remote sensing image retrieval. ISPRS journal photogrammetry remote sensing 145, 197–209 (2018).

18. Cheng, G., Han, J. & Lu, X. Remote sensing image scene classification: Benchmark and state of the art. Proceedings IEEE 105, 1865–1883 (2017).

19. Sumbul, G., Charfuelan, M., Demir, B. & Markl, V. Bigearthnet: A large-scale benchmark archive for remote sensing image understanding. In IGARSS 2019-2019 IEEE International Geoscience and Remote Sensing Symposium, 5901–5904 (IEEE, 2019).

20. Townshend, J. R. & Justice, C. O. Towards operational monitoring of terrestrial systems by moderate-resolution remote sensing. Remote Sensing Environment 83, 351–359 (2002).

21. Morisette, J., Privette, J., Strahler, A., Mayaux, P. & Justice, C. An approach for the validation of global land cover products through the committee on earth observing satellites (2003).

22. McCallum, I., Obersteiner, M., Nilsson, S. & Shvidenko, A. A spatial comparison of four satellite derived 1 km global land cover datasets. International Journal Applied Earth Observation Geoinformation 8, 246–255 (2006).

23. Gao, Y. et al. Consistency analysis and accuracy assessment of three global 30-m land-cover products over the european union using the lucas dataset. Remote Sensing 12, 3479 (2020).

24. Liu, L. et al. Finer-resolution mapping of global land cover: Recent developments, consistency analysis, and prospects. Journal Remote Sensing 2021 (2021).

25. Gengler, S. & Bogaert, P. Combining land cover products using a minimum divergence and a bayesian data fusion approach. International Journal Geographical Information Science 32, 806–826 (2018).

26. Xu, P., Herold, M., Tsendbazar, N.-E. & Clevers, J. G. Towards a comprehensive and consistent global aquatic land cover characterization framework addressing multiple user needs. Remote Sensing Environment 250, 112034 (2020).

27. Fritz, S. et al. Cropland for sub-saharan africa: A synergistic approach using five land cover data sets. Geophysical Research Letters 38 (2011).

28. Zhu, X. X. et al. Deep learning in remote sensing: A comprehensive review and list of resources. IEEE Geoscience Remote Sensing Magazine 5, 8–36 (2017).

29. Shrestha, A. & Mahmood, A. Review of deep learning algorithms and architectures. IEEE Access 7, 53040–53065 (2019).

30. Ma, L. et al. Deep learning in remote sensing applications: A meta-analysis and review. ISPRS journal photogrammetry remote sensing 152, 166–177 (2019).

31. Benhammou, Y., Achchab, B., Herrera, F. & Tabik, S. Breakhis based breast cancer automatic diagnosis using deep learning: Taxonomy, survey and insights. Neurocomputing 375, 9–24 (2020).

32. Rawat, W. & Wang, Z. Deep convolutional neural networks for image classification: A comprehensive review. Neural computation 29, 2352–2449 (2017).

33. Nogueira, K., Penatti, O. A. & Dos Santos, J. A. Towards better exploiting convolutional neural networks for remote sensing scene classification. Pattern Recognition 61, 539–556 (2017).

34. Zhang, L., Xia, G.-S., Wu, T., Lin, L. & Tai, X. C. Deep learning for remote sensing image understanding (2016).

35. Luengo, J., García-Gil, D., Ramírez-Gallego, S., García, S. & Herrera, F. Big data preprocessing - enabling smart data. Cham: Springer (2020).

36. Ghorbanian, A. et al. Improved land cover map of iran using sentinel imagery within google earth engine and a novel automatic workflow for land cover classification using migrated training samples. ISPRS Journal Photogrammetry Remote Sensing 167, 276–288 (2020).

37. Nass, U. Usda-national agricultural statistics service, cropland data layer. United States Department Agriculture, National Agricultural Statistics Service, Marketing Information Services Office, Washington, DC [Available at http://nassgeodata.gmu.edu/Crop-Scape, Last accessed September 2012.] (2003).

38. Yang, L. et al. A new generation of the united states national land cover database: Requirements, research priorities, design, and implementation strategies. ISPRS Journal Photogrammetry Remote Sensing 146, 108–123 (2018).

39. Helber, P., Bischke, B., Dengel, A. & Borth, D. Eurosat: A novel dataset and deep learning benchmark for land use and land cover classification. IEEE Journal Selected Topics Applied Earth Observations Remote Sensing 12, 2217–2226 (2019).

40. Gorelick, N. et al. Google earth engine: Planetary-scale geospatial analysis for everyone. Remote sensing Environment 202, 18–27 (2017).

41. Kennedy, C. M., Oakleaf, J. R., Theobald, D. M., Baruch-Mordo, S. & Kiesecker, J. Managing the middle: A shift in conservation priorities based on the global human modification gradient. Global Change Biology 25, 811–826 (2019).

42. Rottensteiner, F. et al. The isprs benchmark on urban object classification and 3d building reconstruction. ISPRS Annals Photogrammetry, Remote Sensing Spatial Information Sciences I-3 (2012), Nr. 1 1, 293–298 (2012).

43. Penatti, O. A., Nogueira, K. & Dos Santos, J. A. Do deep features generalize from everyday objects to remote sensing and aerial scenes domains? In Proceedings of the IEEE conference on computer vision and pattern recognition workshops, 44–51 (2015).

44. Basu, S. et al. Deepsat: a learning framework for satellite imagery. In Proceedings of the 23rd SIGSPATIAL international conference on advances in geographic information systems, 1–10 (2015).

45. Yang, Y. & Newsam, S. Bag-of-visual-words and spatial extensions for land-use classification. In Proceedings of the 18th SIGSPATIAL international conference on advances in geographic information systems, 270–279 (2010).

46. Dai, D. & Yang, W. Satellite image classification via two-layer sparse coding with biased image representation. IEEE Geoscience Remote Sensing Letters 8, 173–176 (2010).

47. Zhao, B., Zhong, Y., Xia, G.-S. & Zhang, L. Dirichlet-derived multiple topic scene classification model for high spatial resolution remote sensing imagery. IEEE Transactions on Geoscience Remote Sensing 54, 2108–2123 (2015).

48. Zou, Q., Ni, L., Zhang, T. & Wang, Q. Deep learning based feature selection for remote sensing scene classification. IEEE Geoscience Remote Sensing Letters 12, 2321–2325 (2015).

49. Xia, G.-S. et al. Aid: A benchmark data set for performance evaluation of aerial scene classification. IEEE Transactions on Geoscience Remote Sensing 55, 3965–3981 (2017).

50. Van Etten, A. et al. The multi-temporal urban development spacenet dataset. In Proceedings of the IEEE/CVF Conference on Computer Vision and Pattern Recognition, 6398–6407 (2021).

51. Sulla-Menashe, D. & Friedl, M. A. User guide to collection 6 modis land cover (mcd12q1 and mcd12c1) product. USGS: Reston, VA, USA 1–18 (2018).

52. Buchhorn, M. et al. Copernicus global land cover layers—collection 2. Remote Sensing 12, 1044 (2020).

53. Sexton, J. O. et al. Global, 30-m resolution continuous fields of tree cover: Landsat-based rescaling of modis vegetation continuous fields with lidar-based estimates of error. International Journal Digital Earth 6, 427–448 (2013).

54. Teluguntla, P. et al. Global cropland area database (gcad) derived from remote sensing in support of food security in the twenty-first century: current achievements and future possibilities. (2015).

55. Shimada, M. et al. New global forest/non-forest maps from alos palsar data (2007–2010). Remote Sensing environment 155, 13–31 (2014).

56. Hansen, M. C. et al. High-resolution global maps of 21st-century forest cover change. science 342, 850–853 (2013).

57. Simard, M., Pinto, N., Fisher, J. B. & Baccini, A. Mapping forest canopy height globally with spaceborne lidar. Journal Geophysical Research: Biogeosciences 116 (2011).

58. Pekel, J.-F., Cottam, A., Gorelick, N. & Belward, A. S. High-resolution mapping of global surface water and its long-term changes. Nature 540, 418–422 (2016).

59. Gong, P. et al. Annual maps of global artificial impervious area (gaia) between 1985 and 2018. Remote Sensing Environment 236, 111510 (2020).

